# Assessment of the new World Health Organization’s dengue classification for predicting severity of illness and level of healthcare required

**DOI:** 10.1101/516229

**Authors:** Balgees A. Ajlan, Maram M. Alafif, Maha M. Alawi, Naeema A. Akbar, Eman K. Aldigs, Tariq A. Madani

## Abstract

The objective of this observational study was to assess the validity of the new dengue classification proposed by the World Health Organization (WHO) in 2009 and to develop pragmatic guidelines for case triage and management. This retrospective study involved 357 laboratory-confirmed cases of dengue infection diagnosed at King Abdulaziz University Hospital, Jeddah, Saudi Arabia over a 4-year period from 2014 to 2017. The sensitivity of the new classification for identifying severe cases was limited (65.0%) but higher than the old one (30 0%). It had a higher sensitivity for identifying patients who needed advanced healthcare compared to the old one (72.0% versus 32.0%, respectively). We propose adding decompensation of chronic diseases and thrombocytopenia-related bleeding to the category of severe dengue in the new classification. This modification improves sensitivity from 72.0% to 97.5% for identifying patients who need advanced healthcare without altering specificity (96.7%). It also improves sensitivity in predicting severe outcomes from 32% to 88.0%. In conclusion, the new classification had a low sensitivity for identifying patients needing advanced care and for predicting morbidity and mortality. We propose to include decompensation of chronic diseases and thrombocytopenia-related bleeding to the category of severe dengue in the new classification to improve the sensitivity of predicting cases requiring advanced care.

**Author summary:** Dengue fever, the most prevalent arthropod-borne viral disease in human, has been conventionally classified into four main categories: non-classical, classical, dengue hemorrhagic fever, and dengue shock syndrome. Several studies reported lack of correlation between the categories of the conventional classification and the disease severity. As a consequence, the World Health organization proposed in 2008 a new classification that divides dengue into two categories: non-severe and severe dengue; the non-severe dengue is further divided into two categories: dengue with warning signs and dengue without warning signs. In this retrospective study we reviewed 357 cases of dengue diagnosed in our institution over a 4-year period to assess the validity of the new dengue classification in order to develop pragmatic guidelines for case triage and management in the Emergency Departments. We found that the sensitivity of the new classification for identifying severe cases was limited even though it had a higher sensitivity for identifying patients who needed advanced healthcare compared to the old one. We propose adding decompensation of chronic diseases and low platelets-related bleeding to the category of severe dengue in the new classification. This modification dramatically improves the sensitivity for identifying patients who need advanced healthcare and the sensitivity to predict severe outcomes.

## Introduction

Dengue fever (DF) is the most prevalent arthropod-borne viral disease in human and one of the major re-emerging communicable diseases. The World Health Organization (WHO) estimates that around 50 million dengue infections occur annually and approximately 2.5 billion of the world’s population live in dengue-endemic areas (1).

Dengue virus is endemic in Saudi Arabia primarily in the western and southern provinces (2, 3). It is a well-recognized cause of seasonal outbreaks in Jeddah and Makkah (4–7). A large epidemic occurred in 2011 in Jeddah and Makkah when 2569 cases were reported, of which 4 died (Ministry of Health, unpublished data). Another large epidemic occurred in the same region in 2013 when 4411 cases including 8 deaths, were reported (6). Dengue was also reported in other regions of Saudi Arabia, including Al-Madinah (2009), and Aseer and Jizan (2013) (2, 7).

Since 1970s, dengue has been conventionally classified into four main categories (Table 1): non-classical DF, classical DF, dengue hemorrhagic fever (DHF), and dengue shock syndrome (DSS). DHF definition requires the presence of four criteria: fever, thrombocytopenia (<100,000 platelets/mm^3^), hemorrhagic manifestations, and plasma leakage manifest as accumulation of fluids in the peritoneal, pleural, or pericardial spaces, lower limb edema, hypoalbuminemia, or hemoconcentration. Several studies reported lack of correlation between the categories of the conventional classification and the disease severity (8–10). Despite high specificity of the DHF category, the sensitivity is unacceptably low in detecting severe cases of dengue that require specialized care and monitoring in a hospital setting (11–13). As a consequence, a global expert consensus meeting at WHO in 2008 accorded on a new classification of DF. The revised classification divides dengue into two categories (Table 2): non-severe and severe dengue (SDF); the non-severe dengue is further divided into two categories: dengue with warning signs (D+W) and dengue without warning signs (D-W). The new classification was developed based on the level of clinical severity to establish management guidelines and to facilitate dengue reporting and surveillance. Warning signs were proposed to facilitate triage and early detection of potentially severe cases that need hospitalization, particularly in primary care settings and during outbreaks.

**Table 1.**
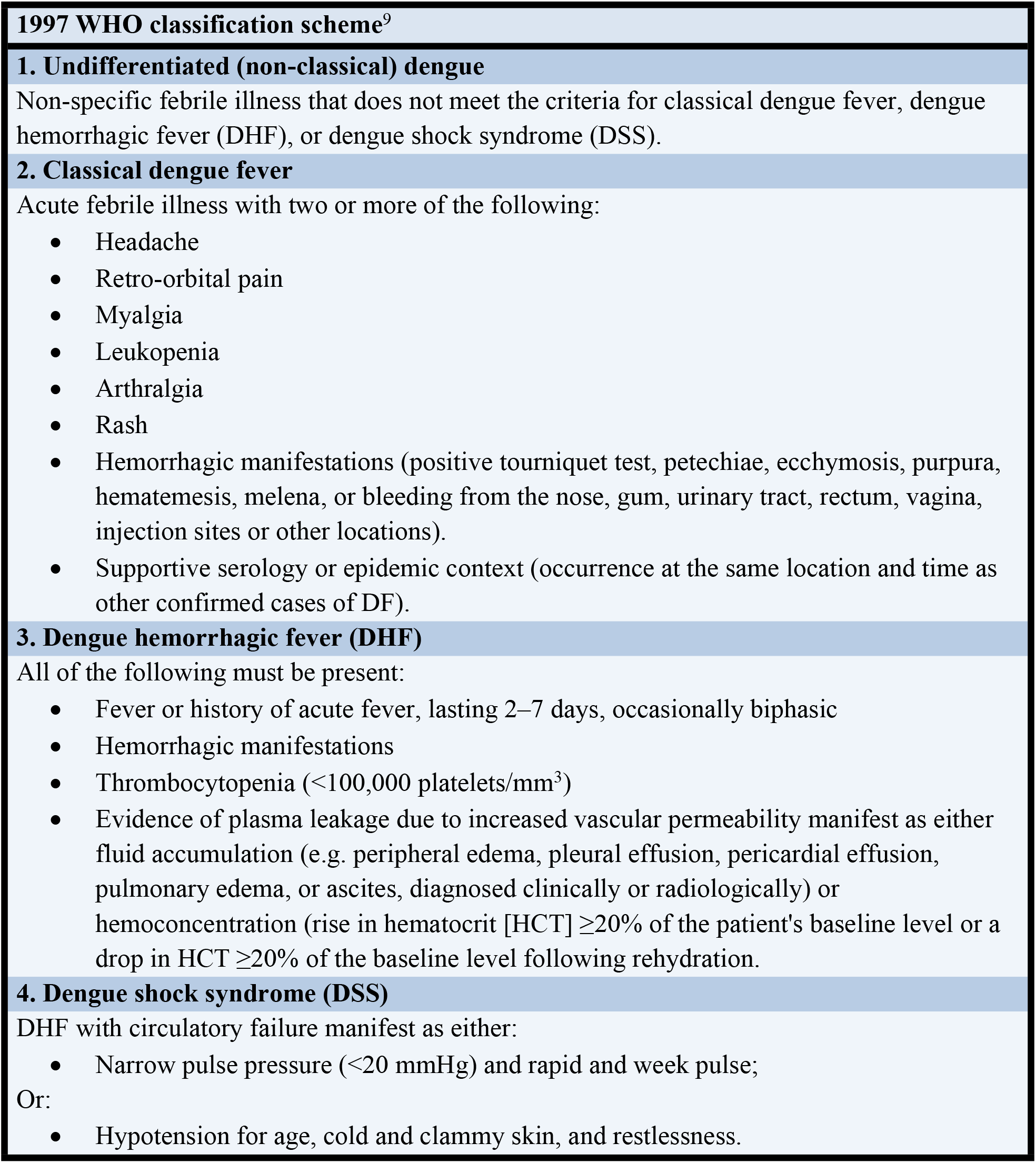
Old World Health Organization (WHO) classifications of dengue (1997)

**Table 2:**
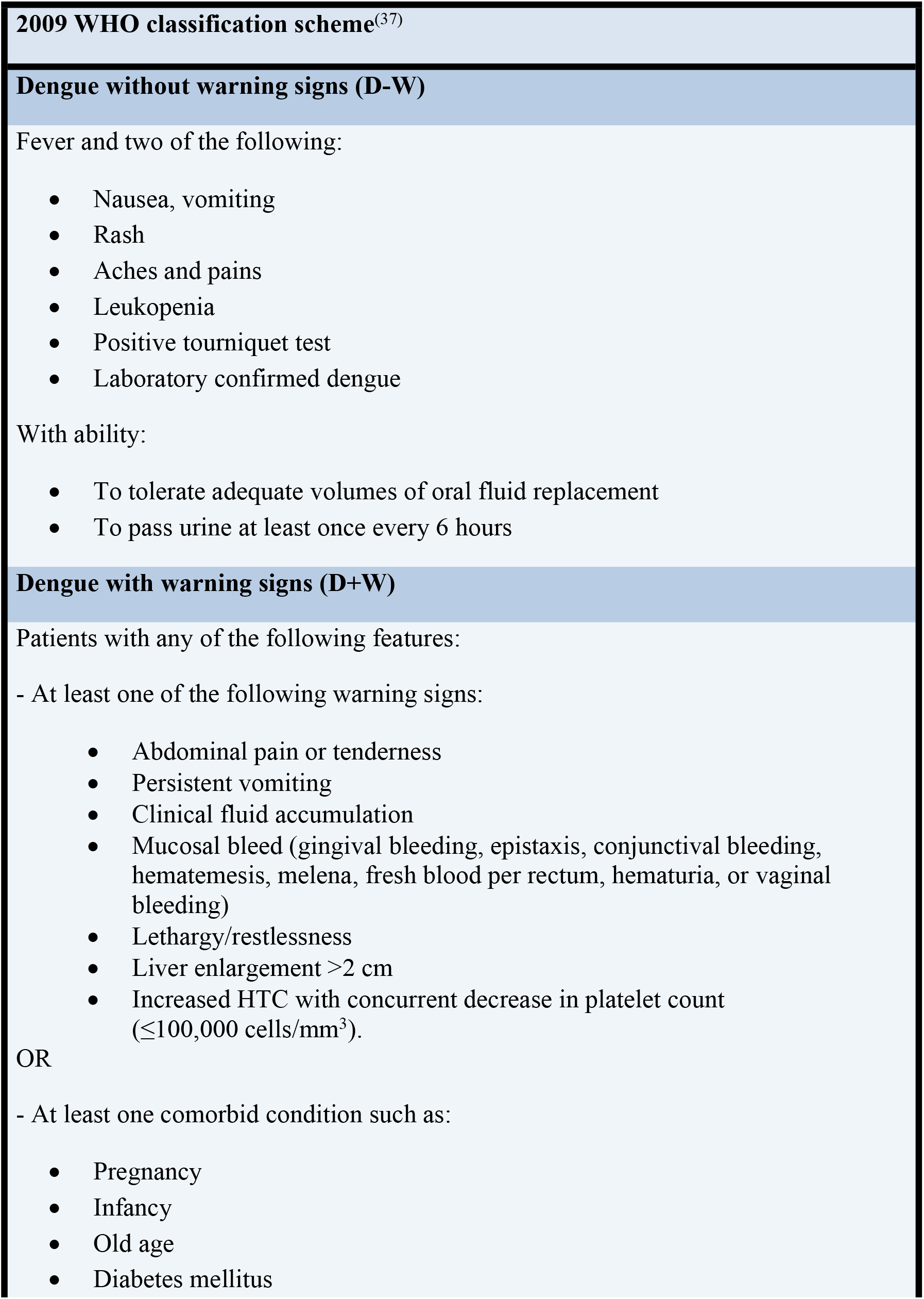
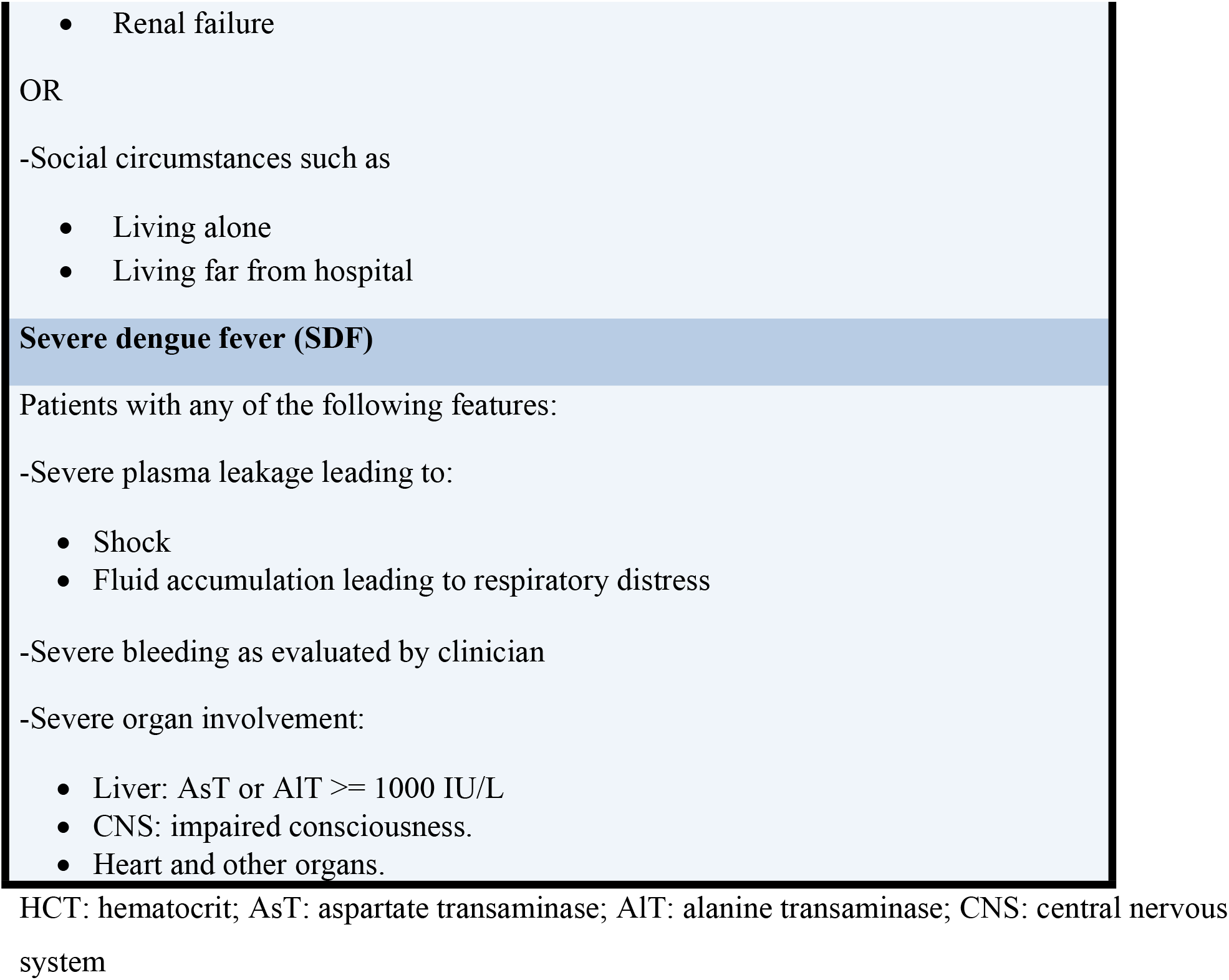
New World Health Organization (WHO) classifications of dengue (2009)

Several studies were conducted to assess the utility of the new dengue classification scheme in clinical practice (14–19). However, current data remain insufficient to establish the validity of this classification in predicting or identifying severe dengue cases that may need close observation or hospitalization for proper management.

This study aimed to assess the validity of the new dengue classification scheme based on data from Jeddah city as part of global evaluation of the new dengue classification.

## Methods

### Population and setting

This retrospective observational study included patients with dengue virus infection reported to the infection control unit at King Abdulaziz University Hospital (KAUH), Western Saudi Arabia over a 4-year period from January 2014 through December 2017. Only laboratory-confirmed cases presenting within 7 days of disease onset were included in the study. The diagnosis was confirmed if at least one of the following criteria was met in acute phase serum: (1) positive reverse transcription polymerase chain reaction (RT-PCR), (2) positive serology for dengue IgM, or (3) positive dengue-specific non-structural antigen-1 (NS1). Exclusion criteria included patients who presented 7 days after the onset of symptoms, or those who were transferred to other hospitals, or whose data were unavailable.

### Definition of cases and warning signs

Persistent vomiting was defined as vomiting at least 5 times per day or vomiting everything the patient ingested. Shock was defined as tachycardia (pulse rate > 100 beats/minute) with either hypotension-for-age or narrow pulse pressure (<20 mmHg). Severe bleeding was defined as major bleeding (hematemesis, melena, or menorrhagia) associated with systolic hypotension, haemoglobin <8 g/dL, or a drop of haemoglobin of >2 g/dL, or bleeding that required blood transfusion. Renal impairment was defined as serum creatinine rise of at least 50% over the baseline that failed to improve after two days of re-hydration. The baseline creatinine was defined as the minimum value following two days of re-hydration. Peak creatinine was defined as the highest creatinine value recorded following two days of re-hydration. The baseline haematocrit (HCT) value was defined as the minimum HCT value following a minimum of two days of re-hydration provided that the patient had passed the 6^th^ day of illness. If no baseline HCT could be defined for a given patient, the hospital’s average value of normal HCT (41% for adults and 42% for pediatrics) was used as the estimated baseline value. Peak HCT was the maximum HCT value recorded during hospital stay. Patients’ clinical outcomes were determined using data from the time of hospital presentation to the time of hospital discharge or demise, and from any subsequent follow up data when available.

Warning signs were only considered at patient’s presentation. The day patients developed severe dengue was documented. Lethargy was not included as a warning sign due to inadequate documentation of this symptom in the patients’ records.

### Patients’ stratification by level of care

Therapeutic interventions and healthcare required by patients were classified into three levels: level I, included patients who were treated on outpatient basis; level II, included hospitalized patients who received intravenous fluids for rehydration and/or those who received platelets due to thrombocytopenia that was not associated with major mucosal bleeding; level III, included hospitalized patients who required intravenous fluids for resuscitation, mechanical ventilation, blood transfusion, inotropic support, or specific treatment for organ failure.

### Statistical analysis

Data were analyzed using IBM SPSS, version 22. P values of <0.05 were considered statistically significant. Chi-square test was used to compare categorical variables. Diagnostic values including sensitivity, specificity, positive predictive value (PPV), and negative predictive value (NPV) of the old and the new WHO classifications for identifying severe cases were analyzed in reference to the level of healthcare the patients required. Thus, the level of healthcare required by patients was the basis for determining severity of dengue in a retrospective manner. For the old classification, patients who were classified as DHF/DSS were considered to have severe dengue, while those classified as non-classical or classical DF were considered to have non-severe dengue. For the new classification, patients classified as SDF were considered to have severe dengue and those classified as dengue with or without warning signs were considered to have non-severe DF. With respect to the gold standard (management level), severe cases were defined as patients who were managed with level III care, whereas non-severe cases were patients who required level I or II care.

### Ethics statement

The Institutional Review Board of King Abdulaziz University Hospital approved the study. Patients’ consent was not required to conduct this retrospective chart review study.

## RESULTS

### Patients’ characteristics

Of 471 laboratory-confirmed dengue cases, 357 met the inclusion criteria. Of those 357 cases, 244 (68·3%) were males; 53 (14·8%) were children (<15 years old) with a mean ± standard deviation (SD) age of 8 ± 4·4 years, and 304 (85·2%) were adults (≥15 years) with a mean age of 33 ± 14.1 years.

The clinical presentation and laboratory characteristics of the 357 eligible patients are summarized in tables 3 and 4. The mean interval from the onset of illness to the hospital presentation was 4 ± 1·8 days (range 1-7, median 4 days).

**Table 3.**
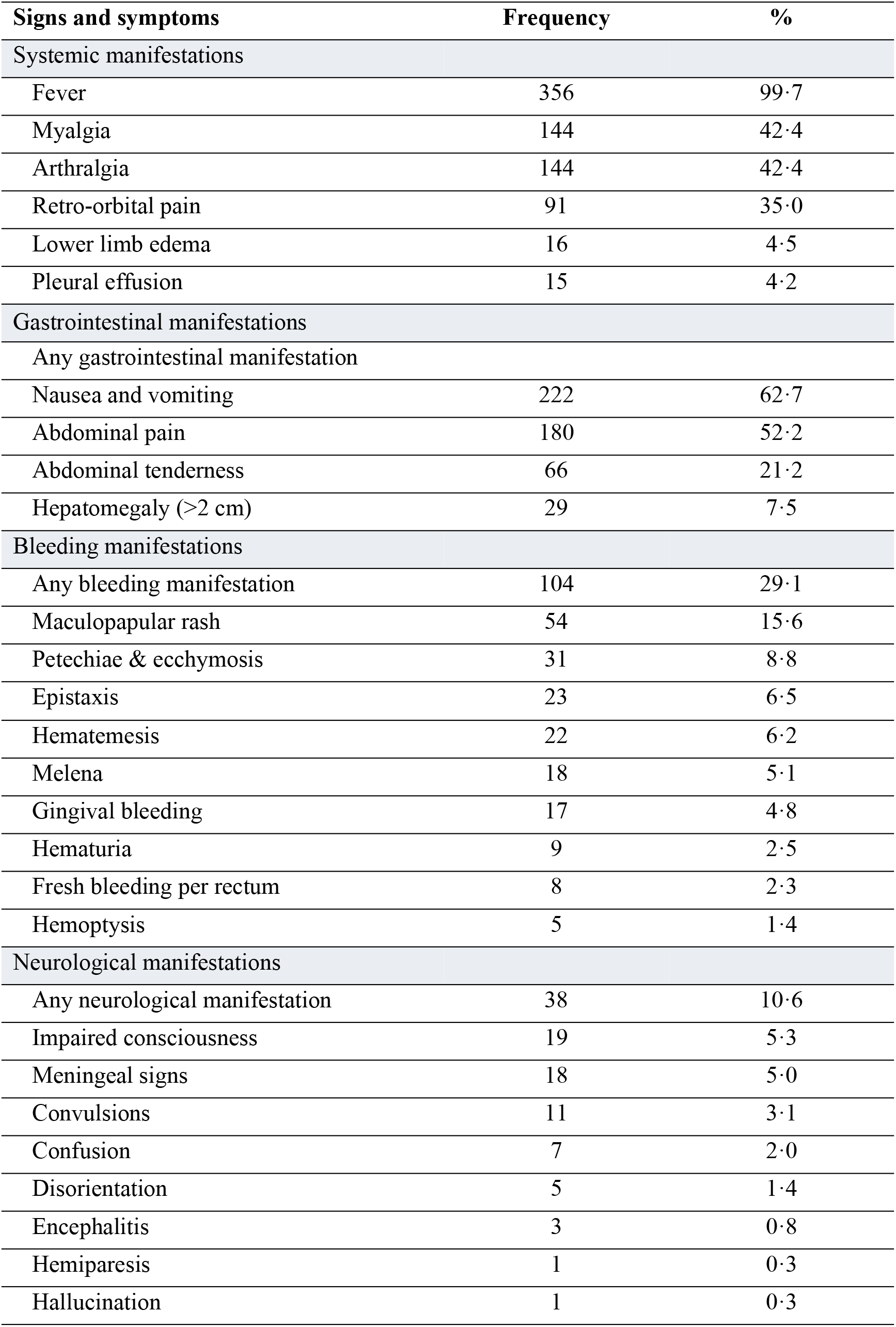
Clinical characteristics of 357 patients with laboratory-confirmed dengue

**Table 4:** Laboratory characteristics of 357 patients with laboratory-confirmed dengue fever

| Parameter | Mean ± standard deviation | Median (range) |
| --- | --- | --- |
| White blood cells ($\times 10^3/\text{mm}^3$) | $3.7 \pm 4.2$ | 2.8 (0.48-66.63) |
| Hemoglobin (g/dL) | $12.3 \pm 2.7$ | 12.7 (1.60-17.80) |
| Maximum hematocrit (%) | $40.9 \pm 6.6$ | 41 (14.7-72) |
| Platelets count at presentation ($\times 10^3/\text{mm}^3$) | $108.4 \pm 85.7$ | 99 (4-765) |
| Lowest platelets count ($\times 10^3/\text{mm}^3$) | $83.8 \pm 67.4$ | 70 (2-396) |
| Albumin (g/L) | $32.7 \pm 6.5$ | 34 (4-62) |
| Highest AsT (IU/L) | $249.3 \pm 741.8$ | 101 (12-7592) |
| Highest ALT (IU/L) | $187.9 \pm 5.64$ | 77 (13-6956) |
| **Confirmatory tests** | **Frequency** | **%** |
| Serology IgM |  |  |
| Positive | 282 | 79.0 |
| Negative | 60 | 16.8 |
| RT-PCR |  |  |
| Positive | 180 | 50.4 |
| Negative | 144 | 40.3 |
| NS1 antigen |  |  |
| Positive | 123 | 34.5 |
| Negative | 96 | 26.9 |

AsT: aspartate transaminase; ALT: alanine transaminase; RT-PCR: Reverse transcription polymerase chain reaction; NS1: nonstructural protein 1.

### Management and level of care

Of 357 eligible patients, 190 (53·2%) patients were hospitalized for a mean of 7·6 ± 9·8 days (range 1-95, median 5 days). Nineteen (5·3%) patients required admission to the intensive care unit (ICU) and 8 (2·2%) patients died. Of 357 eligible patients, 167 (46·8%) were assessed and managed in the emergency/ambulatory departments.

### Evaluation of the classifications in screening for level III of care

Cross-tabulations of the level of care (level I/II versus level III) with the old and the new classifications showed 86% and 93% proportional agreement (PA) of the old and the new classifications with the level of care, respectively (Tables 5 and 6).

**Table 5:**
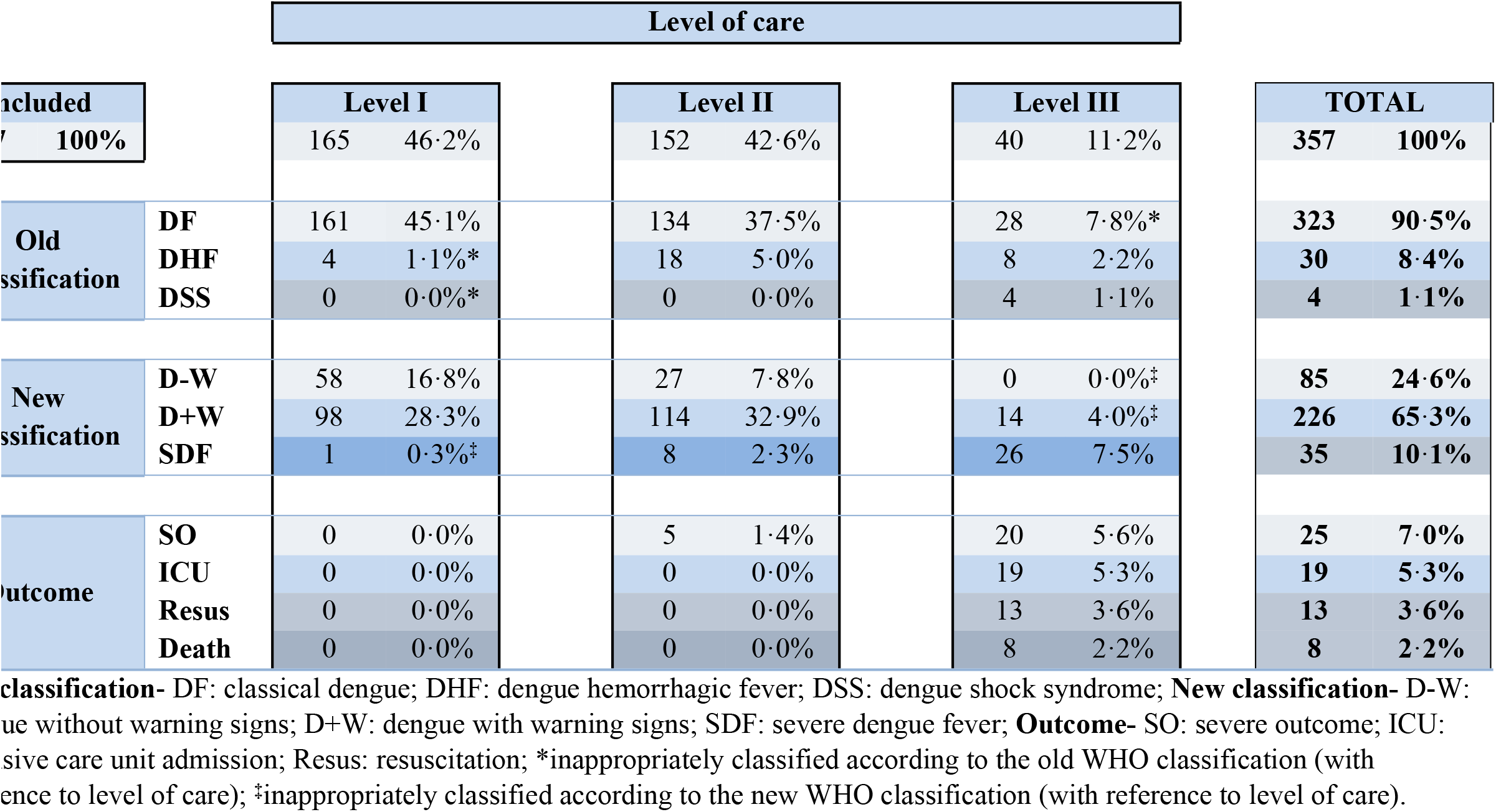
Correlation of dengue fever classifications on admission with actual level of care required

**Table 6:** Two-by-two contingency table of the old and the new WHO classifications with the level of care

| **Old WHO classification (test)** | **Level of care (N = 357)** |  | **Total** |
| --- | --- | --- | --- |
|  | **Level I and II** | **Level III** |  |
| DF [negative] | 295 (82·6); [TN] | 28 (7·8); [FN] | **323 (90·5)** |
| DHF/DSS [positive] | 22 (6·2%); [FP] | 12 (3·4); [TP] | **34 (9·5)** |
| **Total** | **317 (89·4)** | **40 (10·6)** | **357 (100·0)** |

TN: true negative; FN: false negative; FP: false positive; TP: true positive. Proportional agreement (95% CI) = 86% (0·82-0·90); Kappa (95% CI) = 0·25 (0·1-0·4); SE = 0·076.

| **New WHO classification (test)** | **Level of care (N = 346)** |  | **Total** |
| --- | --- | --- | --- |
|  | **Level I and II** | **Level III** |  |
| D-W/D+W [negative] | 297 (85·8); [TN] | 14 (4·9); [FN] | **311 (89·9)** |
| SDF [positive] | 9 (2·6); [FP] | 26 (7·5); [TP] | **35 (10·1)** |
| **Total** | **306 (88·4)** | **40 (11·6)** | **346 (100·0)** |

TN: true negative; FN: false negative; FP: false positive; TP: true positive. Proportional agreement (95% CI) = 93% (0·93-0·96); Kappa (95% CI) = 0·66 (0·53-0·79); SE = 0·07.
Results are presented as number (%). DF: dengue fever; DHF: dengue hemorrhagic fever; DSS: dengue shock syndrome; D-W: dengue without warning signs; D+W: dengue with warning signs; SDF: severe dengue fever.

There was a minimal agreement (Kappa standard error [SE] = 0.179 [0 075]; p=0 01) between the two classifications, although proportional agreement (PA) was 85·3% (p=0.003) (Table 6).

Diagnostic values including sensitivity, specificity, PPV and NPV of the old and the new classifications in identifying patients who required level III of care is presented in table 7.

**Table 7:**
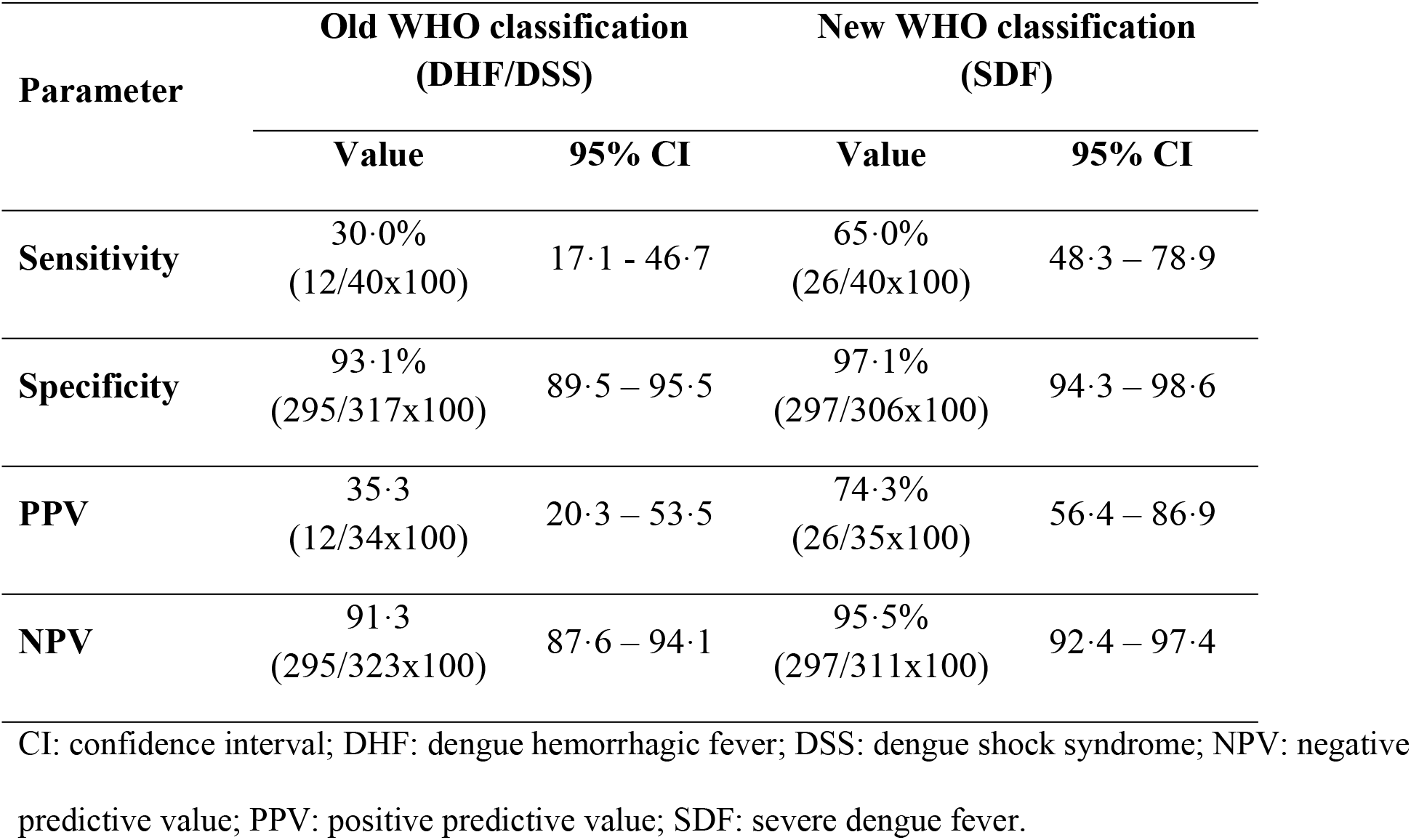
Diagnostic values of the old and the new WHO dengue fever classifications in detecting patients requiring level III of care

| Parameter | Old WHO classification<br>(DHF/DSS) |  | New WHO classification<br>(SDF) |  |
| --- | --- | --- | --- | --- |
|  | Value | 95% CI | Value | 95% CI |
| **Sensitivity** | 30.0%<br>(12/40x100) | 17.1 - 46.7 | 65.0%<br>(26/40x100) | 48.3 – 78.9 |
| **Specificity** | 93.1%<br>(295/317x100) | 89.5 – 95.5 | 97.1%<br>(297/306x100) | 94.3 – 98.6 |
| **PPV** | 35.3<br>(12/34x100) | 20.3 – 53.5 | 74.3%<br>(26/35x100) | 56.4 – 86.9 |
| **NPV** | 91.3<br>(295/323x100) | 87.6 – 94.1 | 95.5%<br>(297/311x100) | 92.4 – 97.4 |

CI: confidence interval; DHF: dengue hemorrhagic fever; DSS: dengue shock syndrome; NPV: negative predictive value; PPV: positive predictive value; SDF: severe dengue fever.

### Proposed revision of new WHO classification

In an attempt to improve the sensitivity of the clinical assessment in identifying patients who would require level III of care, we propose two revised versions of the new classification by integrating new criteria to the definition of SDF. The first proposed revision integrates patients with acutely decompensated chronic disease; e.g. patients with known cardiomyopathy who present with acute heart failure or patients with known hematological disease, such as sickle cell anemia, presenting with aplastic crisis. This first revision improves the sensitivity from 65·0% to 85·0%, the PPV from 74·3% to 79·1%, and the NPV from 95·5% to 98·0%, with no change in specificity (971%). The second proposed revision also integrates (besides decompensated chronic disease) patients with thrombocytopenia of <20,000 platelets/mm^3^ with any bleeding. This second revision results in further improvement of the sensitivity from 65·0% to 97·5%, the PPV from 74·3% to 79·6%, and the NPV from 95·5% to 99·7%, with insignificant reduction of the specificity from 97·1% to 96·7%. The diagnostic value of the first and second proposed revisions of SDF in the new classification is presented in table 8 and the revised criteria are presented in table 9.

**Table 8:**
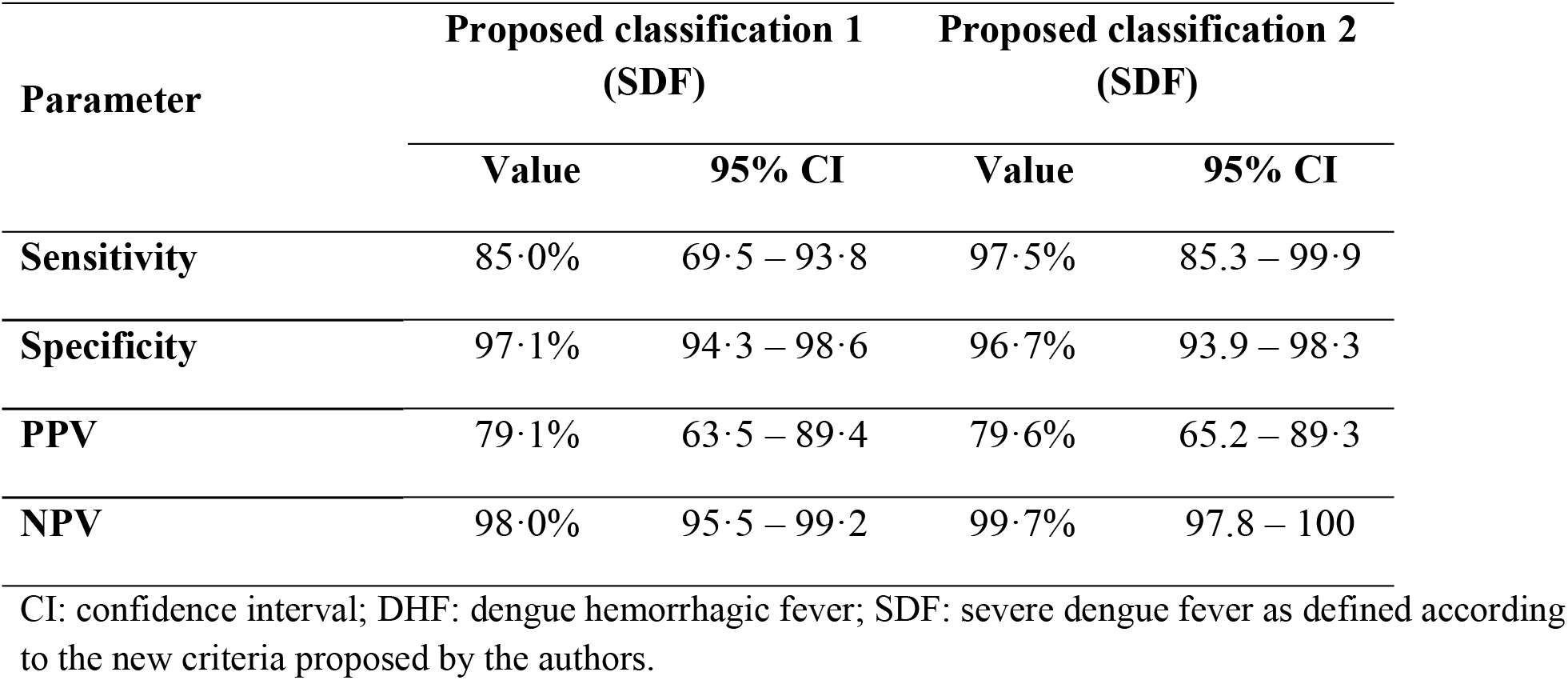
Diagnostic values of two proposed modifications (1 and 2) of the new WHO dengue fever classification in identifying patients with severe dengue requiring level III of healthcare

**Table 9.** Definition of severe dengue fever (SDF) in the new World Health Organization (WHO) classification and two proposed revisions:

| **Original criteria for diagnosing severe dengue fever** | **1<sup>st</sup> proposed revision** | **2<sup>nd</sup> proposed revision** |
| --- | --- | --- |
| Patients with any of the following features: | Patients with any of the following features: | Patients with any of the following features: |
| **A-** Severe plasma leakage leading to: | **A-** Severe plasma leakage leading to: | **A-** Severe plasma leakage leading to: |
| 1- Shock or | 1- Shock or | 1- Shock or |
| 2- Fluid accumulation leading to respiratory distress | 2- Fluid accumulation leading to respiratory distress | 2- Fluid accumulation leading to respiratory distress |
| **B-** Severe bleeding as evaluated by clinician | **B-** Severe bleeding as evaluated by clinician | **B-** Severe bleeding as evaluated by clinician |
| **C-** Severe organ involvement: | **C-** Severe organ involvement: | **C-** Severe organ involvement: |
| • Liver: AsT or ALT ≥ 1000 IU/L<br>• CNS: impaired consciousness<br>• Heart and other organs | • Liver: AsT or ALT ≥ 1000 IU/L<br>• CNS: impaired consciousness<br>• Heart and other organs | • Liver: AsT or ALT ≥ 1000 IU/L<br>• CNS: impaired consciousness<br>• Heart and other organs |
|  | **D-** Acute decompensation of chronic disease | **D-** Acute decompensation of chronic disease |
| | | **E-** Minor bleeding with thrombocytopenia $<20,000$ platelets/mm<sup>3</sup> |

### Screening for severe outcomes

Severe outcomes were defined as patients who recovered after having complications directly or indirectly related to dengue or those who died. The sensitivity of the new classification, the 1^st^ proposed revision, and the 2^nd^ proposed revision, for identifying patients with severe outcomes was 72·0%, 84·0%, and 88·0%, respectively, and the NPV was 97·7%, 98·7% and 99·0%, respectively (Table 10).

**Table 10:** Diagnostic values of the old, the new, and the two proposed modifications (1^st^ and 2^nd^) in identifying patients with severe outcomes

|  | Old WHO classification |  | New WHO classification (original) |  | 1<sup>st</sup> proposed revision |  | 2<sup>nd</sup> proposed revision |  |
| --- | --- | --- | --- | --- | --- | --- | --- | --- |
|  | Value | 95% CI | Value | 95% CI | Value | 95% CI | Value | 95% CI |
| **Sensitivity** | 32·0% | 15·7 – 53·6 | 72·0% | 50·4 – 87·1 | 84·0% | 63·1 – 94·7 | 88·0% | 67·7 – 96·8 |
| **Specificity** | 92·2% | 88·6 – 94·7 | 94·7% | 91·5 – 96·8 | 93·1% | 89·6 – 95·6 | 91·6% | 87·9 – 94·3 |
| **PPV** | 23·5% | 11·4 – 41·6 | 51·4% | 34·3 – 68·3 | 48·8% | 33·6 – 64·3 | 44·9% | 30·9 – 59·7 |
| **NPV** | 94·7% | 91·5 – 96·9 | 97·7% | 95·2 – 99·0 | 98·7% | 96·4 – 99·6 | 99·0% | 96·8 – 99·7 |

NPV: negative predictive value; PPV: positive predictive value.

### Warning signs as predictors of level of care and outcome severity

Evaluation of the warning signs of the new classification showed that patients with hemoconcentration associated with concurrent drop in platelets count had approximately 5-fold increased risk of requiring level III healthcare (OR [95% CI] = 4·97 [2·35–10·50], p<0 001) and approximately 7-fold increased risk of severe outcome (OR [95% CI] = 6·89 [2·90-16·37], p<0 001). Other warning signs were not significant predictors of level III healthcare or severe outcome (Table 11). Correlation between the number of warning signs (<2 versus ≥2), the old and the new classifications, and the level of care is presented in table 12.

**Table 11:** Warning signs as predictors for level of care and severe outcome (univariate regression analysis)

| Warning sign (predictor) | Dependent variable |  |  |  |  |  |  |  |
| --- | --- | --- | --- | --- | --- | --- | --- | --- |
|  | Level III of care |  |  |  | Severe outcome |  |  |  |
|  | OR | 95% CI |  | p-value | OR | 95% CI |  | p-value |
| Persistent vomiting | 2.05 | 0.91 | 4.6 | 0.085 | 0.53 | 0.12 | 2.31 | 0.396 |
| Abdominal pain | 0.70 | 0.35 | 1.38 | 0.301 | 0.76 | 0.33 | 1.72 | 0.507 |
| Abdominal tenderness | 0.86 | 0.34 | 2.17 | 0.752 | 2.22 | 0.92 | 5.40 | 0.077 |
| Clinical fluid accumulation | 1.27 | 0.36 | 4.50 | 0.710 | 2.25 | 0.62 | 8.18 | 0.220 |
| Mucosal bleeding | 1.73 | 0.82 | 3.67 | 0.152 | 0.80 | 0.27 | 2.41 | 0.798 |
| Hepatomegaly >2 cm | 0.33 | 0.04 | 2.53 | 0.288 | 3.70 | 1.26 | 10.85 | 0.017* |
| Laboratory warning signs<sup>#</sup> | 4.97 | 2.35 | 10.50 | <0.001* | 6.89 | 2.90 | 16.37 | <0.001* |

\*statistically significant result ( $p < 0.05$ ).
<sup>#</sup>Laboratory warning signs: hemoconcentration with concurrent drop in the platelets count.
Abbreviations, OR: odds-ratio; CI: confidence interval.
Multivariate regression model including hepatomegaly and lab warning signs as predictors of severe outcome showed OR [95% CI] = 2.53 [0.80 – 8.04], $p = 0.115$ and 6.14 [2.53 – 14.88], $p < 0.001$ , respectively.

**Table 12:** Correlation between the number of clinical warning signs (<2 versus ≥2) in the old and the new dengue classifications and the required level of care

| Parameter | Category | < 2 warning signs<br>(N=246) |  | ≥2 warning signs<br>(N=111) |  | p-value |
| --- | --- | --- | --- | --- | --- | --- |
|  |  | Freq. | % | Freq. | % |  |
| **Old classification** | DF | 237 | 96·3 | 86 | 77·5 | <0·00001* |
|  | DHF/DSS | 9 | 3·7 | 25 | 22·5 |  |
| **New classification** | D-W/D+W | 215 | 90·3 | 96 | 88·9 | 0·679 |
|  | SDF | 23 | 9·7 | 12 | 11·1 |  |
| **Level of care** | Level I/Level II | 222 | 90·2 | 95 | 85·6 | 0·196 |
|  | Level III | 24 | 9·8 | 16 | 14·4 |  |

DF: dengue fever; DHF: dengue hemorrhagic fever; DSS: dengue shock syndrome; D-W: dengue without warning signs; D+W: dengue with warning signs; SDF: severe dengue fever; because of missing data, frequencies do not sum up to totals in new classification.

## DISCUSSION

This study compared the old and the new WHO dengue classifications as predictors of both levels of healthcare and patients’ outcomes among 357 patients with confirmed dengue. It demonstrated that both classifications were inadequate in identifying patients who required advanced level of healthcare. Both classifications had a low-to-moderate sensitivity (30·0% to 65·0%), although the new classification had an acceptable sensitivity in predicting severe outcomes (72·0%). Consequently, we propose to revise the new classification by integrating 2 new criteria in the definition of SDF, namely, acute decompensation of chronic diseases and evidence of minor bleeding in association with thrombocytopenia <20,000 platelets/mm^3^. The proposed revision improved the sensitivity to 97·5% in detecting patients who need level III healthcare without altering specificity, which remained high (96·7%). It also improved sensitivity of predicting severe outcomes to 88·0%. Furthermore, the proposed revisions had 99·7% and 99·0% NPVs for the level of care and outcome severity, respectively. This means that a patient who would not be classified as SDF according to our proposed criteria would have 0·3% probability to require level III of healthcare and 1·0% probability to have severe outcome including complications and mortality; versus 4·5% and 5·3% for the original classification, respectively.

Data from literature generally suggest that the new classification had a higher sensitivity as well as a higher or comparable specificity in identifying severe cases requiring higher level of healthcare in comparison with the old classification (14–19). Several studies that compared the WHO classifications showed that a significant proportion of patients with SDF were misclassified as classical DF using the old classification, while they were readily identified by using SDF in the new classification (14–16). Other studies compared the accuracy of the two classifications in certain clinical contexts. For example, a Brazilian study of 267 pediatric cases, showed that the old classification had a lower sensitivity (62·3% versus 86·8%) but a higher specificity (93·4% versus 73·0%) in discriminating severe cases compared to the new classification (20). Another Brazilian retrospective study evaluating 121 autopsied individuals who died during 2011-2012 dengue epidemics, showed that the new classification had a higher sensitivity to discriminate dengue deaths and that the absence of plasma leakage and thrombocytopenia were the main reasons for failure of the old classification to discriminate DHF cases (21). A review by Bandyopadhyay et al., analyzed 37 studies using the old WHO dengue classification, and demonstrated that this classification had a low sensitivity with frequent overlap between the different classes, especially in endemic areas (22). Additionally, the new classification has been shown to be a better measure for case reporting and surveillance (21–23). Conversely, other authors found no difference between the two classifications in term of sensitivity (24, 25).

The new classification intended to develop a system that helps in directing patients’ management and improving clinical outcome by reducing morbidity and mortality. Beyond the controversy over the usefulness of the old and the new WHO definition of SDF in identifying severe cases, the practicability of the new definition in epidemic contexts was contested, as it was perceived to be entailing heavier workload to healthcare personnel (24). According to the new classification, all patients presenting with dengue warning signs should be admitted for observation. This would raise the hospitalization rate from 9.5% (DHF/DSS) to 75.4% (D+W/SDF) according to our data, resulting in unnecessary observation/admission, which might overwhelm healthcare personnel and resources particularly during outbreaks. As shown in our study, almost half of D+W patients were managed as outpatients, which makes the relevance of warning signs in terms of disease severity questionable (14–17). Only hemoconcentration with concurrent drop in the platelet count was a significant predictor of severe outcome and the need for advanced healthcare. Most of the previous studies found that none of the suggested warning signs was a significant predictor of dengue severity (24, 26, 27), although one of them showed that the presence of five or more warning signs was a significant predictor of SDF (27). A multicenter study across 18 countries, assessing user-friendliness and acceptance of the new classification from health professionals’ viewpoint, showed that 24·1% of health professionals had concerns with the new classification including: a possible increase of hospitalization rates, non-specificity of warning signs, a possible increase of cost if more patients were admitted, and the need for more training and dissemination of more concise clinical protocols (17). These disadvantages called for a revision to improve its practicability and specificity in identifying severe cases.

In the present study, the new definition of SDF missed 14 (35%) patients who needed advanced (level III) medical care. Of these 14 cases, 8 had co-existing hematological conditions (sickle cell anemia and hereditary spherocytosis) that caused severe drop in hemoglobin level necessitating urgent blood transfusion, 5 patients had severe thrombocytopenia (<20,000 cells/mm^3^) with evidence of bleeding, and one patient had a neurological deficit that was exacerbated by dengue. In addition, 12/35 patients with severe outcome (morbidity or mortality) had thrombocytopenia <50,000 cells/mm^3^ at presentation, 2 of them were <20,000 cells/mm^3^. Adding two new criteria, namely, thrombocytopenia <20,0000 cells/mm^3^ with evidence of bleeding, even though minor, and decompensated chronic illness, to the definition of SDF improved its sensitivity to identify patients who needed advanced level of care from 65·0% to 97·5% and its sensitivity to predict cases likely to have morbidity or mortality from 72·0% to 88·0%.

The first criterion suggested to be added to our proposed revision of the new classification as an indication of SDF is “acute decompensation of a preexisting comorbidity”. Previous studies also showed that pre-existence of a comorbid condition was strongly associated with morbidity and mortality in DF (28). A systematic literature review by Toledo et al, analyzed 16 studies from Taiwan, Singapore, Brazil, Pakistan, India, and Thailand investigating clinical risk factors of severe dengue, beyond the WHO classification criteria. The review demonstrated that patients with cardiovascular, metabolic, renal, or respiratory comorbidities, as well as older patients were at higher risk for developing severe form of dengue regardless of whether they fulfill severity criteria according to the new WHO classification (29). Another study from Puerto Rico focusing on dengue severity in the elderly, showed that elderly patients were at increased risk for hospitalization and death, although they were less likely than the young patients to present with hemorrhagic manifestations (30). Another case-control study from Singapore compared 818 cases classified as DHF with 1467 cases classified as DF and found that diabetes mellitus, hypertension, and age ≥60 years were associated with a higher risk for developing DHF (odds ratios: 1·78, 2·16, and 3·44, respectively) (31). These observations indicate that comorbidities and older age, not only increase the likelihood of developing DHF and DSS, but also contribute to the clinical severity and outcome in the absence of classifiable severity criteria. Therefore, the new classification should be revised to include comorbidities and risk stratified by age.

The second criterion suggested to be included in our proposed revision of the new SDF definition is “thrombocytopenia≤20,000 platelets/mm^3^ with any bleeding, even though minor”. Thrombocytopenia <100,000 cells/mm^3^ constitutes only one of the warning signs in the original version of the new WHO classification (2009) but it is not included in the definition of SDF. In a French-Polynesian study, thrombocytopenia <20,000 platelets/mm^3^ was associated with a longer hospital stay, more frequent admission to ICUs, and higher mortality (32). However, only two thirds of cases were classified as DHF/DSS and the remaining one third was classified as DF resulting in underestimation of the severity of illness (32). This is consistent with our results demonstrating that severe thrombocytopenia (<20,000 platelets/mm^3^) is a strong indicator for severe dengue and the need for advanced level of care. Therefore, the new classification should be revised to include severe thrombocytopenia in the definition of SDF to improve identification of severe cases.

In our study, 9/35 patients classified as SDF received level II of healthcare and the remaining 26 patients were hospitalized for close observation and monitoring. Of these 26 patients, 3 patients had suspicion of liver damage (aminotransferases >1000 U/L), which proved to be a transient elevation of liver enzymes resolving spontaneously without complications. Among the patients who had neurological manifestations, impaired consciousness was the most frequent sign (5·3%), followed by meningeal signs (5·0%), and convulsion (3·1%) with one patient presenting with encephalitis. All cases with neurological manifestations benefited from conservative management. Similar to our findings, a Vietnamese study reported that impaired consciousness was the most frequent neurological manifestation (33). Impaired consciousness is part of SDF criteria according to the new classification. A study by Gupta et al, demonstrated a greater risk for neurological complications among patients classified as classical DF in the old classification which did not include neurological symptoms or low level of consciousness at presentation as signs of SDF (33). Recent reviews also highlighted a rising trend of neurological complications of dengue and that it is associated with more frequent hemorrhagic manifestations, higher prevalence of DSS, and increased hospital stay and mortality (34). A study from Brazil reported 21·2% of neurological manifestations with confusion being the most frequent sign (35). In Europe, an even higher prevalence (24·0%) of neurological manifestations was reported among imported cases of dengue infection, mainly including cases acquired in Asia and the Americas (36). This variability in the prevalence of neurological manifestations may be explained by varying time of clinical assessment with respect to patient’s first presentation and symptoms onset. Furthermore, lack of clear definitions of organ damage that reflects the actual disease severity may lead to misplacement of patients under SDF category. In our cohort, only 60·9% of patients having organ damage needed level III care. Thus, standardized definition of organ damage is needed to improve the classification specificity.

Limitations of our study include the retrospective design yielding incomplete observations, especially for cases treated on outpatient basis; which compromised the power of analysis. Moreover, most of the severe cases presented at a late stage, which limited the accuracy of warning signs assessment and analysis as predictors for severe dengue.

### Conclusion

The new WHO dengue classification had low sensitivity for identifying patients in need of advanced level of care and for predicting morbidity and mortality. This classification needs to be revised to improve its sensitivity in predicting the required level of care. It is proposed to include 2 additional criteria to the definition of SDF, namely, decompensation of chronic disease and thrombocytopenia <20,000 cells/mm^3^ with evidence of any bleeding, even though minor. Adding these two criteria substantially improves the sensitivity to predict cases requiring advanced level of care and to predict severe outcomes. Further studies and meta-analyses are needed to support the validity of the proposed modification in different settings. Further, criteria such as organ damage and impaired consciousness should be distinctively defined to avoid overlap of the definitions and misclassification.

## Acknowledgments

We thank Fatmah Hassan Ba-Obaid, Najla Mubarak Alharbi, Rana Abdulrahman Alamoudi, and Daniya Osama Abdouh for their assistance in data collection.

## Supporting Information Legends

STROBE Checklist

